# Early Detection of Tau Pathology

**DOI:** 10.1101/2021.05.14.444233

**Authors:** Parag Parekh, Andrew Badachhape, Qingshan Mu, Rohan Bhavane, Mayank Srivastava, Igor Stupin, Prajwal Bhandari, Laxman Devkota, Jason Eriksen, Eric Tanifum, Ketan Ghaghada, Ananth Annapragada

## Abstract

While a definitive Alzheimer’s disease (AD) diagnosis remains a post-mortem exercise, the ATN Research Framework proposed by the National Institute on Aging and the Alzheimer’s Association utilizes a score representing the presence of amyloid deposits (A), tau deposits (T) and neuronal degeneration markers (N), with A+T+ necessary for a positive diagnosis. Current detection of tau pathology lags amyloid detection by years and by the time both markers are detected the disease is fairly advanced. We describe the development of a new generation of molecular imaging probes for *in vivo* detection of cells undergoing abnormal phosphorylation representing the initial stages of pTau pathology, potentially enabling a very early stage diagnosis of AD. We describe a novel nanoparticle formulation that binds such abnormally phosphorylating cells in a mouse model of tau pathology, enabling in *vivo* visualization of the hyperphosphorylative state by magnetic resonance imaging. Our results demonstrate the potential of this novel platform to identify a correlative marker signifying the development of future tau pathology, and has implications for early-stage diagnosis of Alzheimer’s disease.

## Introduction

The microtubule associated protein tau, coded for by the *MAPT* gene, is abundant in the brain and is present in neurons, glia and other cell types. Tau shows immense diversity. It is expressed in six isoforms and has a vast array of post-translational modifications, including glycosylation, glycation, nitration, ubiquitination and more than 80 phosphorylation sites expanding the complexity of its role in health and disease^1, 2^. A definitive feature of many neurodegenerative diseases including AD, frontotemporal lobar degeneration (FTLD), Parkinson’s disease (PD) is the presence of intracellular aggregated filamentous tau (collectively termed “tauopathies”)^3–6^. This pathogenic insoluble tau been implicated in disease progression in these dysfunctional brains. The transition from physiological soluble tau is primarily associated with changes in its phosphorylation state leading to oligomeric tau^7^, then tau fibrils known as paired helical fragments (PHF) that collectively form characteristic neurofibrillary tangles (NFT). The tau aggregates are then capable of “infecting” a healthy cell inducing further misfolding, aggregation, and neuro-toxicity^8, 9^. Studies of intercellular propagation demonstrate passage through an extracellular phase that progresses throughout the brain ^10, 11^.

The research framework proposed for precise diagnosis of AD by the National Institute of Aging-Alzheimer’s Association (NIA-AA) categorizes extracellular deposits of amyloid beta (A), presence of intraneuronal hyperphosphorylated tau (T) and markers of neurodegeneration or neuronal injury (N). Each biomarker is scored either positive or negative^12^. To be on the AD continuum, A+ (Amyloid positive) is required, while a positive diagnosis of AD requires A+ and T+. Biomarker detection can be by i) positron emission tomography (PET) imaging of amyloid and tau or, ii) cerebrospinal fluid (CSF) detection of reduced Aβ_42_, and/or high Aβ_40_/ Aβ_42_ and high phosphorylated tau and total tau, or iii) neuronal injury or degeneration as shown by structural brain magnetic resonance imaging (MRI)^13^. Tracking of brain pathology in longitudinal studies suggests that tau pathology may precede the accumulation of Aβ, but is undetectable as it is below current biomarker detection threshold levels, and is amplified catastrophically by independent Aβ deposition ^14–16^. ATN research framework based diagnosis of AD is therefore limited by tau pathology detection, providing an impetus to develop specific and sensitive tau detection methods for earlier diagnosis.

Other factors to consider in the development of tau detection methods include the invasive nature of CSF sampling requiring lumbar puncture, and in the case of PET imaging, exposure to ionizing radiation, high cost, well documented side effects, irregular availability in primary care setting, and uneven geographical availability of PET scanners and isotopes^17–21^. The short half-life of PET agents also poses challenges for the detection of intracellular tau in the early stages of tau pathology formation^22^. Blood based markers are very promising, but only provide an indirect measure that cannot provide information on the localization of tau pathology in brain^23–26^. Methods to detect early tau pathology that avoid these pitfalls are therefore highly desirable.

The initiation of tau pathology is marked by abnormal phosphorylation of tau^27, 28^. We hypothesize that hyperphosphorylative conditions in neurons, consistent with an altered balance of kinase-phosphatase activity resulting in elevated levels of hyperphosphorylated tau species, result in unique surface markers^29–31^. We envisioned a targeted contrast-enhanced MRI test identifying the earliest stages of tau pathology represented by cells in the process of abnormally phosphorylating tau. We used an iterative Cell-SELEX process to identify DNA thioaptamers that specifically bound such cells^32, 33^. The use of phosphorothioate modified aptamers enhances stability under physiological conditions. High T1 relaxivity PEGylated liposomes bearing macrocyclic Gd-chelates^34^ were modified to bear the aptamers on their surface, thus enabling targeting of the particles to the surface of hyperphosphorylative cells for contrast-enhanced MRI. At the molecular level, we sought the binding targets of the aptamers, identifying vimentin, a normally intracellular protein that interestingly is specifically expressed on the surface of cells under hyperphosphorylative conditions, and representing a possible biomarker of pathological hyperphosphorylation found in AD.

## Results

### Validation of cell surface changes in hyperphosphorylative conditions

SH-SY5Y, a human neuroblastoma cell line that can be differentiated into neuron-like cells by changes in culture medium was used to model cell-surface changes under hyperphosphorylative conditions (Fig 1B). We used retinoic acid (RA) to induce cell differentiation marked by temporal changes in morphology including the formation and lengthening of neurites and with a strong increase in levels of intracellular tau. Imbalance in the kinase and phosphatase activity leading to hyperphosphorylation, simulating early stages of tauopathies, was induced by the use of a cell permeable neurotoxin okadaic acid (30nM, 24 h) (OA)^35^, and confirmed by the increase in phosphorylated tau S202/T205 and S396 (Supp. Fig. 1). In parallel experiments, a milder agent, excitotoxin quinolinic acid (QA) 1µM for 24 h was also used to induce hyperphosphosphorylation^36^.

**Figure 1:**
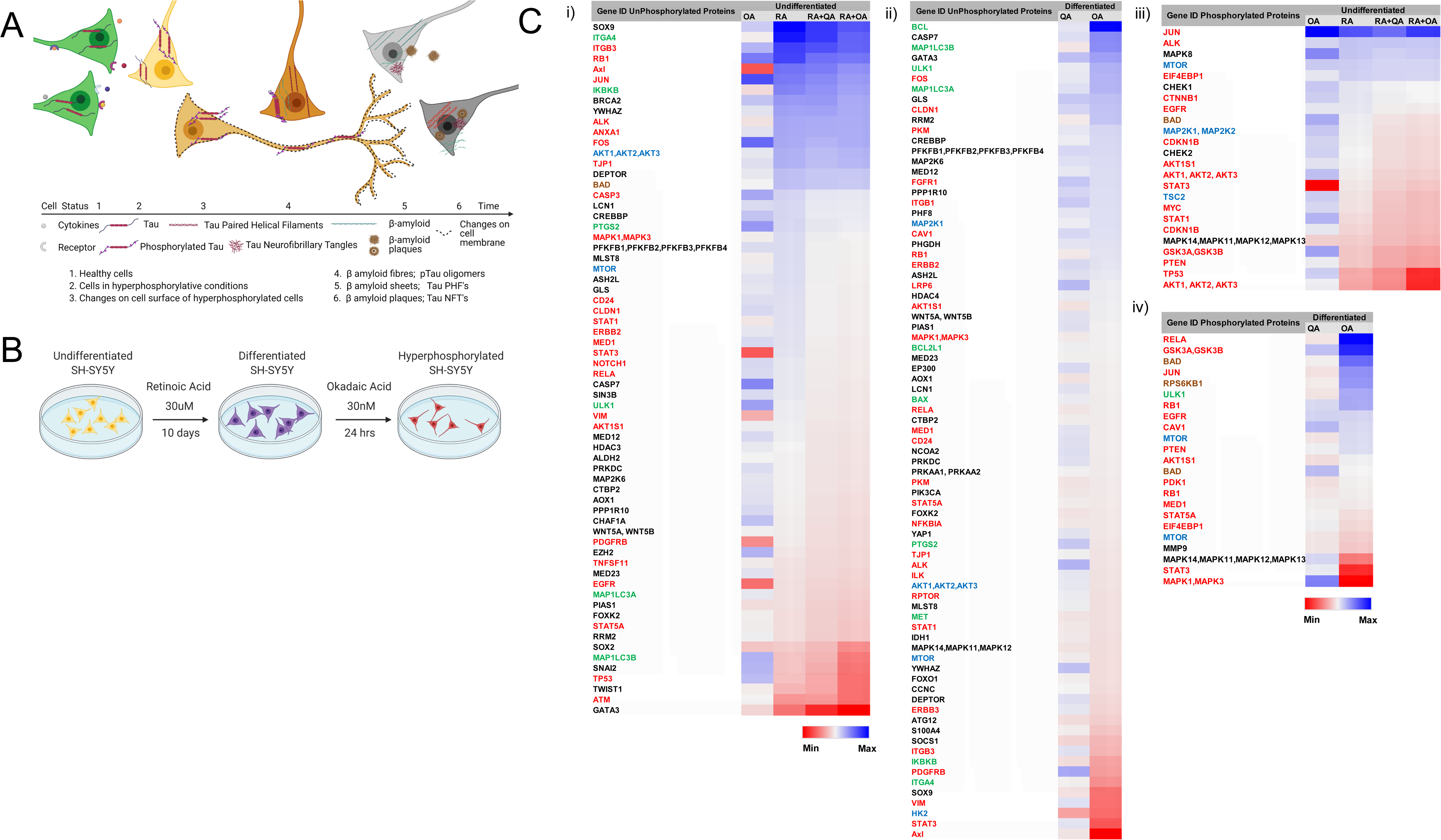
Hyperphosphorylation and cell surface changes in a cell of neuronal phenotype. **(A)** *<u>Cell changes during hyperphosphorylation</u>*: Tau hyperphosphorylation is a precursor to tau fibrillation. We hypothesize that cell surface changes occur in parallel with tau hyperphosphorylation. These changes may constitute a marker for early detection of AD. **(B)** *<u>In</u> <u>vitro model of hyperphosphorylation in a cell of neuronal phenotype</u>*: Differentiated SH-SY5Y cells are used to model hyperphosphorylation. Hyperphosphorylation can be induced by either okadaic acid (OA) or quinolinic acid (QA) **(C)** *<u>Protein analysis at different stages of</u> <u>hyperphosphorylation</u>*: Reverse phase protein array analysis of undifferentiated, retinoic acid (RA) differentiated and hyperphosphorylative cells (by Okadaic acid OA or Quinolinic acid QA). Heat map depicts changes in total protein expression. Proteins expressed on cell-membrane are depicted in red, peripheral –membrane proteins in blue and single-pass and multipass membrane proteins in green. Panel i) depicts changes in un-phosphorylated proteins on undifferentiated and ii) differentiated SH-SY5Y cells. Changes due to phosphorylation treatments with OA and QA on differentiated cells are shown in panels iii) undifferentiated and iv) differentiated SH-SY5Y cells

A reverse phase protein array (RPPA) assay was conducted to test the hypothesis that hyperphosphorylation results in compositional changes reflected on the cell-surface. RPPA analysis demonstrated marked cell surface changes in hyperphosphorylative cells including over-expression of cell surface receptors. RPPA assessment with a panel of 221 proteins across different pathways and post-translational modifications including phosphorylation examined SH SY5Y cell lysates under differentiated and hyperphosphorylative states. We identified 98 proteins and 36 phosphorylated proteins that showed significant change in their expression under different conditions (**Fig 1C**). Uniprot ^37^ protein associations showed 44 cell-membrane associated proteins, 10 peripheral membrane or 12 single-pass membrane proteins were significantly altered under hyperphosphorylative conditions.

### Screening for aptamers that bind cells in hyperphosphorylative state

Aptamer screening was performed using the cell-SELEX approach on differentiated SH-SY5Y cells in a hyperphosphorylative state (Fig. 2A). A total of 26 cell SELEX cycles were performed. To remove oligonucleotides that bound common cell surface molecules not specific to the hyperphosphorylative state, a negative selection was introduced at cycles 12 and 13 using differentiated, non hyperphosphorylative cells (i.e. without OA treatment). Anticipating that selected aptamers would be systemically delivered as nanoparticle imaging agents, and the primary toxicity driven by anomalous hepatocyte uptake, we conducted another round of negative selection at cycles 20 and 21 using a hepatocyte cell-line THLE-3 to remove oligonucleotides that exhibited enhanced uptake by hepatocytes.

**Figure 2:**
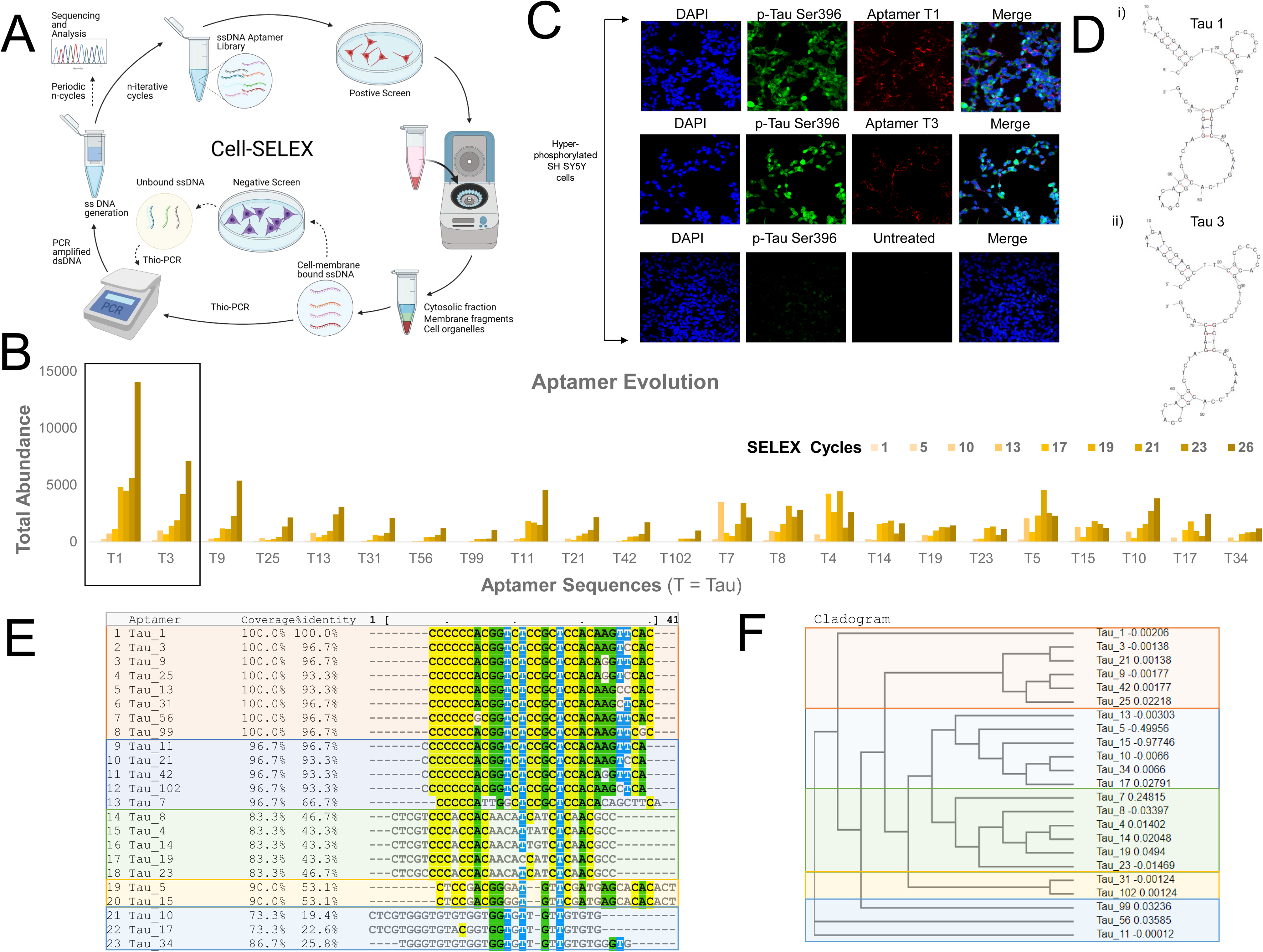
Cell-SELEX to identify biomarkers onset of AD. Okadaic acid treated differentiated SH-SY5Y cells were used as a surrogate for hyperphosphorylative neurons to screen DNA aptamers that specifically recognize the differences between the surfaces of treated and untreated cells, using the cell-SELEX methodology modified to capture membrane binding aptamers. (A) Pictorial representation of the cell-SELEX process (B) Abundance of the top 23 sequences from SELEX cycle 1 - 26 depicting their evolution. Note that the fractions are low until about cycle 10, when they increase sharply. Note that the abundance of Tau 1, Tau 3 continually increase with increasing cycle number. (C) Hyperphosphorylated SH-SY5Y cells stained with 50nM Cy5 labelled Tau1, Tau3 aptamer and untreated for 2h at 4⁰C.(D) i) Tau 1 and ii) Tau 3 secondary structure using Mfold. (E) Multiple sequence alignment of the top 23 aptamer sequences by MAFFT. Note the existence of five families. (F) Cladograms showing relationship between the Tau aptamer sequences and the aptamer families.

### Tau1 and Tau3 aptamers specifically bind hyperphosphorylative cells

Sequencing of all the selected pools was performed using the Ion Torrent sequencing platform^38^ and revealed the evolution of families of DNA sequences, with enrichment particularly evident after 10 rounds of SELEX. Negative selection eliminated certain sequences that were not specific to the hyperphosphorylative state, or promoting hepatocyte uptake. However, the relative abundance of key sequences increased steadily throughout the whole process. The 23 most abundant sequences at round 26 were identified and their abundance throughout the SELEX process as calculated using AptaAligner^39^ is shown in Fig. 2B . The sequence Tau1 exemplifies this behavior representing a whole 59% of cycle 26. A single base difference from this sequence, Tau3, is the second most represented sequence. The sequences present at the final round were grouped by hierarchical clustering and sequence homology using the multiple sequence alignment code MAFFT ^40^ showing five distinct families (Fig 2E) also presented as a cladogram (using the Clustal Omega^41^) showing the common ancestry between these five aptamers families (Fig. 2F). Elevated levels of Tau1 and Tau3 binding to the membrane of hyperphosphorylative cells is demonstrated in Figure 2C. The secondary structure of the aptamers Tau1 and Tau3 calculated using mfold^42^ is shown in Fig 2D. The apparent equilibrium dissociation constants (Kd _app_) were measured by serial dilution of aptamer solutions with target hyperphosphorylated and non-target differentiated and undifferentiated SH-SY5Y cells. The affinity of these aptamers was also tested with another immortal neural progenitor stem cell line ReN-VM^43^ in hyperphosphorylative and non-hyperphosphorylative conditions . The Kd _app_ for Tau1 and Tau3 with hyperphosphorylated SH SY5Y cells is 0.167 ± 0.015 nM and 0.194 ± 0.032 nM; and for the ReN-VM cells 318.15 ± 46.2 nM and 234.24 ± 38.6 nM respectively (Supp. Fig 2).

### TauX nanoparticles for magnetic resonance imaging

For *in vivo* imaging of the hyperphosphorylative state in the brain of live mice, aptamer-targeted nanoparticles were fabricated for use as a molecular MRI contrast agent (TauX). TauX, was formulated as two versions, one using the Tau1 aptamer (TauT1) and another using the Tau3 aptamer (TauT3). Liposomal nanoparticles were synthesized using a lipid mixture that included lipidized Gd-DOTA for MR imaging, cholesterol for liposomal stability; additionally we also incorporated lipidized rhodamine for studying *ex-vivo* microscopic distribution of liposomal nanoparticles in brain tissues using fluorescence microscopy^44, 45^. The TauX compositions also included DSPE-mPEG2000 to increase the *in vivo* circulation time^34^. Particles had a hydrodynamic diameter of ∼150nm, ∼86,000 Gd-chelates per liposomes and ∼500 aptamers conjugated to the outer leaflet of each liposomal nanoparticle (Fig.3)

**Figure 3:**
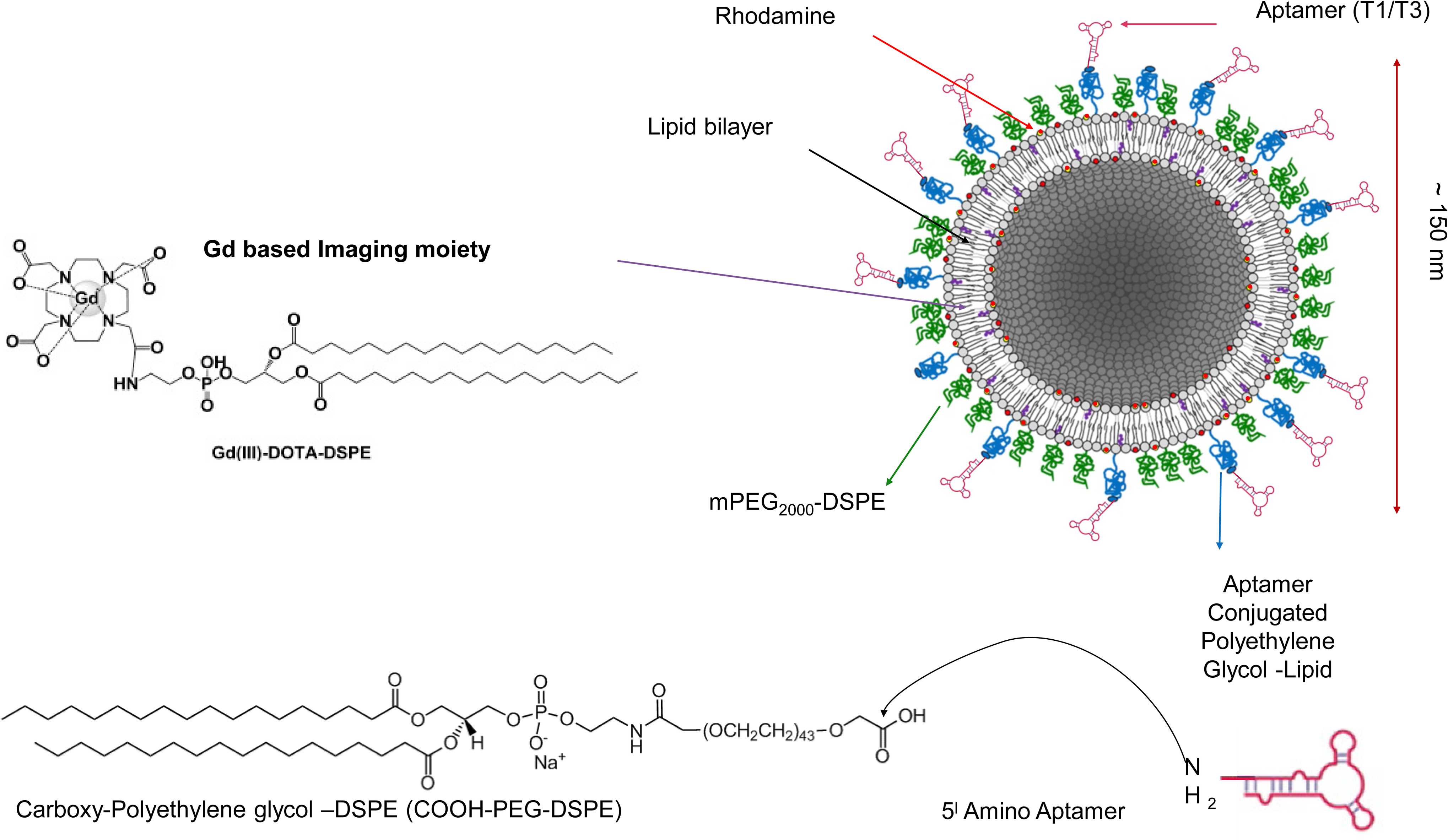
Aptamer-targeted liposomal-Gd nanoparticle TauX contrast agent. The liposomal bilayer incorporates DSPE-DOTA-Gd for MR contrast, DSPE-PEG2000 to enhance circulation half-life, DSPE-PEG3400-aptamer (Tau1/Tau3) for targeting and lissamine rhodamine for fluorescence imaging.

### *In vivo* molecular MRI using TauX for detection of hyperphosphorylative cells

To test if TauX could detect the hyperphosphorylative state *in vivo*, MRI studies were performed in a P301S transgenic mouse model of AD-related tauopathy. Studies were performed in transgenic and age-matched wild type mice at 2-3 months of age. At this young age, transgenic animals do not show frank tau pathology (i.e., neurofibrillary tangles), but practically all will develop tau pathology by around 8 months of age. Animals underwent baseline, pre-contrast MRI. Thereafter, animals were intravenously administered MRI contrast agent (TauT1, TauT3 or non-targeted control stealth liposomes that were not expected to provide signal enhancement as they did not have a targeting aptamer). Delayed post-contrast MRI was performed 4 days later. MR images were acquired using a T1-weighted spin-echo (T1w-SE) sequence and a T1-weighted fast spin echo inversion recovery (FSE-IR) sequence^46^. Transgenic mice administered TauT1 and TauT3 demonstrated signal enhancement in the cortex and the hippocampus regions of the brain (**Fig. 4B**). Wild-type mice (WT) administered TauT1 or TauT3 did not show signal enhancement in the brain. Similarly, transgenic mice administered non-targeted liposomal-Gd contrast agent did not show signal enhancement in cortex or hippocampus. These regions of interest were further analyzed quantitatively and signal-enhancement between the transgenic and wild-type mice were found to be statistically significant (p < 0.05) (Fig 4D). A baseline enhancement threshold of ∼6% (=2X standard deviation of signal in baseline scans) was used as the classification threshold. Animals that showed signal enhancement above the threshold were identified as positives. Receiver operator characteristic (ROC) curves were generated with a six-point ordinal scale to assess sensitivity and specificity for detecting the genotype, using TauT1 and TauT3 contrast agents, and constructed over the entire tested group, including controls. The aptamer-targeted nanoparticle contrast agents, TauT1 and TauT3, showed overall AUC and accuracy of ∼0.95. TauT3 demonstrated higher sensitivity than TauT1.

**Figure 4:**
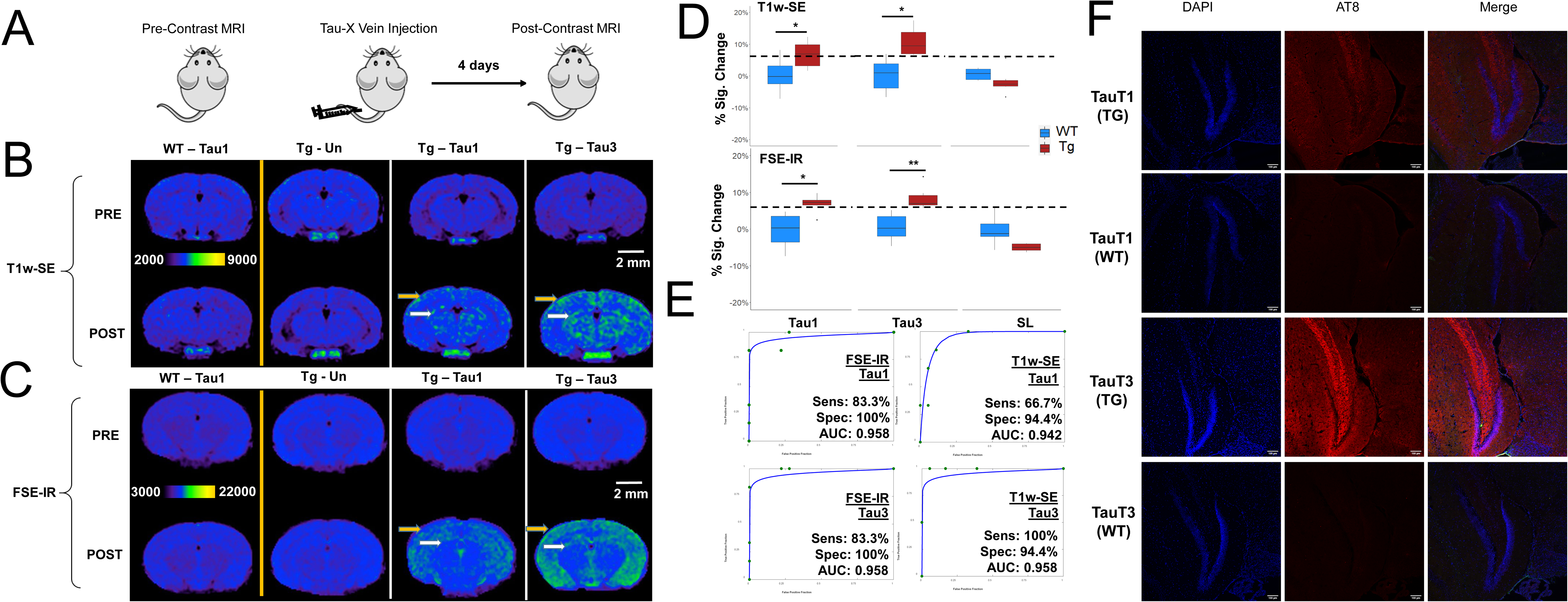
T1-weighted spin-echo (T1w-SE) and (FSE-IR) images demonstrate signal enhancement as TauX binds to hyperphosphorylated tau *in vivo*. **(A)** In vivo imaging was performed on a 1 T permanent MRI scanner (M7, Aspect Imaging). Two month old mice were injected with 0.2mM Gd dose TauX NP. Pre- and post-contrast images were acquired with the following parameters: Imaging sequence: T1 weighted spin echo (T1w-SE) TR = 600 ms, TE = 11.5 ms, slice thickness = 1.2 mm, matrix = 192x192, FOV = 30 mm, slices = 16, NEX = 4. Animals were euthanized after final imaging session and brains harvested for post-mortem analysis. (**B**) and (**C**) demonstrate signal enhancement in delayed post-contrast scans of transgenic (Tg) P301S mice treated with TauX relative to age-matched controls. Wild type (WT) animal shows no signal enhancement four days after administration of TauX. Tg animal showed high enhancement in cortical (yellow arrow) and hippocampal regions (white arrow) four days after administration of TauX. Transgenic animal shows no signal enhancement four days after injection of untargeted contrast (UC). Scale bar represents 2 mm. All animals are shown on the same color scale. (**D**) Signal enhancement in TG animals relative to WT counterparts and UC-treated Tg animals for both T1w-SE and FSE-IR sequences (*p<0.05 ; **p<0.005). (**E**) The Receiver operating characteristic (ROC) curve demonstrates TauX accuracy in identifying early age Tg animals. ROC curve is generated by fitting observed operating points for true positive fraction versus false positive fraction (green dots) using a six point ordinal scale. A fitted curve (blue) connects the observed operating points. Area under curve (AUC) is calculated using the fitted curve. Sensitivity describes the true positive rate for Tg mice given TauX (n=6) while specificity is the true negative rate for WT mice given TauX (n=6), WT mice given UC Gd nanoparticle contrast (n=6), and Tg mice receiving UC Gd nanoparticle contrast (n=6). (**F**) The mice selected for TauX injection were genotyped which only revealed the presence of transgene. We confirmed the presence of hyperphosphorylated tau species in the mice by immunofluoscence on the brains sections harvested post MRI scans. Brains were perfused with heparin-PBS and fixed in OCT. IF with antibody AT8 probed for presence of ptau species provided high concordance with genotype confirming the presence of hyperphosphorylative conditions in the mice.

Post-mortem brain analysis was performed in 2-3 month old transgenic and wild-type mice. Immunofluorescence analysis using AT8 antibody staining revealed the presence of hyperphosphorylated tau species in transgenic mice but absent in wild type mice (**Fig. 4F**). A 100% concordance was observed between AT8 positivity and animal genotype. In summary, *in vivo* studies demonstrated that TauX enabled *in vivo* molecular MRI of the hyperphosphorylative state months before frank tau pathology i.e. the presence of neurofibrillary tangles, becomes evident in transgenic mice.

### Target identification of aptamers

To characterize the binding target of the aptamers, we performed an aptamer-based pulldown assay, aptamer-based immunoprecipitation, followed by mass-spectrometry. We performed the assay for both Tau1 and Tau3, aptamers. A ranking of the abundance scores for identified proteins revealed keratin 6a, Keratin 6b and Vimentin as possible binding targets (Table 1). The surface expression of Keratin 6a, 6b was similar on wild-type and transgenic tissue sections whereas the Vim expression was higher in the transgenic mice (Fig 5C). SH-SY5Y cells under undifferentiated, and differentiated hyperphosphorylative conditions show increasing levels of vimentin (Fig.5B) further suggesting it is a potential target of aptamers Tau1 and Tau3.

**Figure 5:**
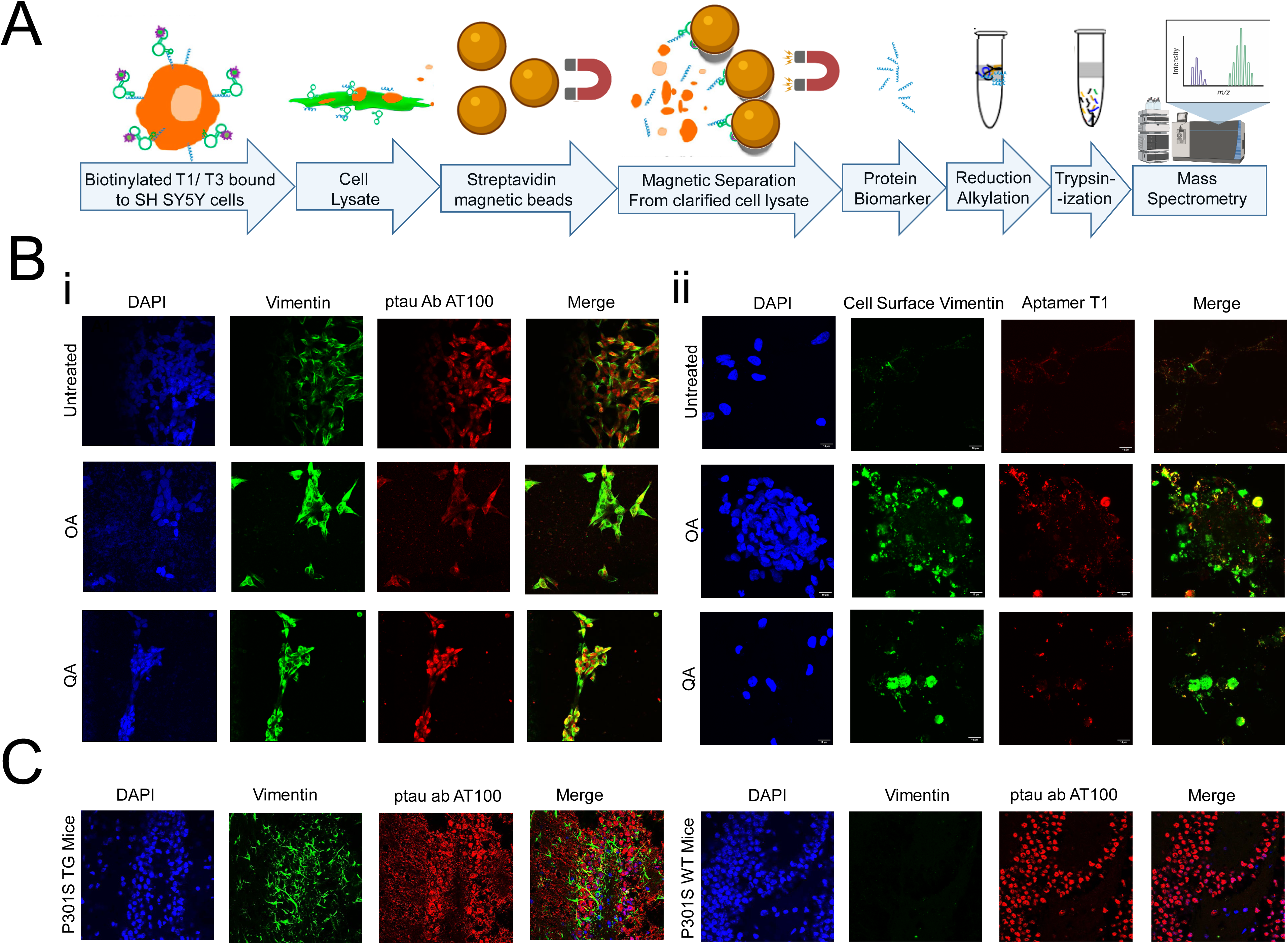
Proteomic analysis for aptamer target identification. A. Procedure for tryptic digest of aptamer bound cell membrane analyzed by LC/MS/MS. The raw data files were processed and searched against the SwissProt 2012 01 (Human) database. Vimentin was identified as possible target. B. Presence of the aptamers binding target on SH SY5Y cells; undifferentiated and hyperphosphorylated OA (24h, 30nM) and QA (24h, 100nM); i) co-stained with Vimentin (D21H3) and ptau (AT100) antibodies or ii) cell surface vimentin (Clone 84-1) antibody and aptamer T1 (50nM); nuclei counterstained with DAPI. C. Expression of Vimentin in P301S TG and WT frozen mouse tissue sections stained with Vimentin (SP20) and ptau (AT100) antibodies and DAPI stained nuclei.

**Figure 6:**
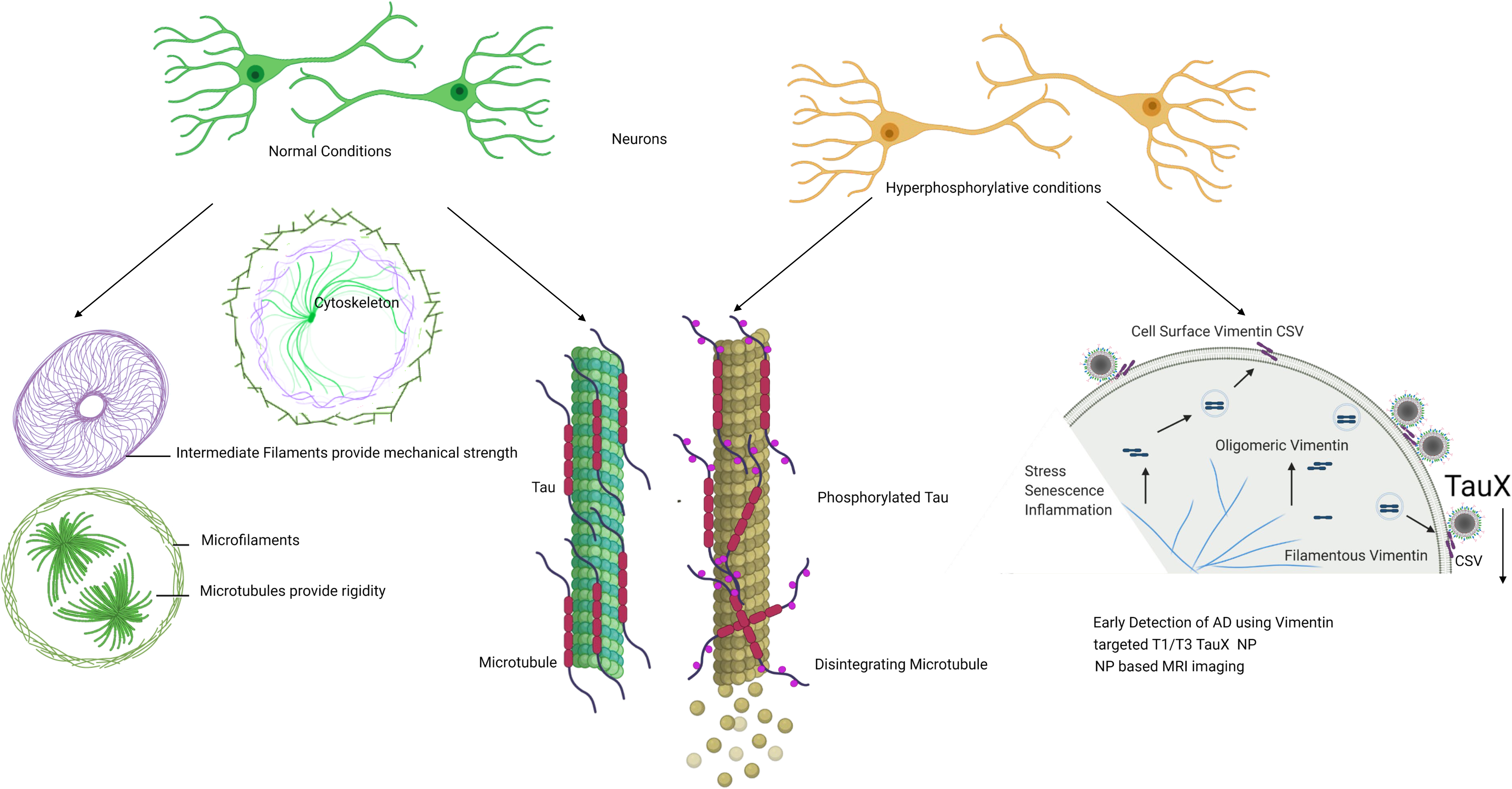
Early detection of future tau pathology. Hyperphosphorylative conditions provoke neuronal cells to undergo changes including on the cell surface. Translocation of Vimentin to the cell surface was exploited to generate an aptamer that was developed for *in vivo* use as targeted liposomal nanoparticle TauX for early MRI detection of tau pathology in a mouse model

## Discussion

The ATN research framework suggests biomarkers to diagnose and classify AD. Under this framework CSF based detection of Aβ, tau (total, and phosphorylated) have been reported but only at the prodromal stage of disease, in patients with mild cognitive impairment^47^. Non-invasive neuroimaging tools, such as structural MRI to diagnose and monitor neurodegeneration^48^ show a definitive correlation with cognitive decline, visualizing atrophic regions that depict neuronal injury in late stage disease^49^. However, a reliable marker of early presymptomatic stage disease is yet to be identified.

While the role of Aβ and tau in the development of AD and the mechanism of transition from presymptomatic to symptomatic AD are yet unclear, the time scale of the transition is generally accepted to be over a period of 10-20 years^14^. Aβ deposits are considered the start of neurodegeneration but recent studies indicate that tau pathology^50^ shows a stronger correlation with disease progression suggesting that the limitation of current tests is their inability to identify early stage pathological tau^15^. CSF presence of hyperphosphorylated tau species p-181 and p-217 is associated with Aβ deposition that precedes a positive tau PET^51^ but only has a concordance of 50%-70% ^52^. Taken as a whole, the roles of Aβ and tau deposition in disease progression and the role of Aβ in the spread of initial tau aggregates, strongly suggest that a biomarker of pathological tau at a presymptomatic stage of the disease is likely to advance detection by several years, and constitutes the motivation for this work.

Initial tau aggregation is thought to be triggered by an imbalance in cellular homeostasis caused by dysregulated phosphorylation^53, 54^. Several kinases can phosphorylate tau at multiple locations; at least 45 sites have been observed experimentally^55–57^. Combined with reduced phosphatase activities in AD, the altered kinase-phosphatase balance yields hyperphosphorylative conditions that cause abnormal hyperphosphorylation of tau^58^. Disruption of the normal function of tau, modulating microtubule dynamics by lowering its binding capabilities^59^ increases the level of cytosolic free tau leading to aggregation and fibrillization of tau that spreads throughout the connected brain, seeding the pathology^60, 61^. We hypothesized that this initial process of hyperphosphorylation is associated with changes on the surface of hyperphosphorylative cells. We therefore sought to identify these surrogate markers of tau hyperphosphorylation that presage future tau pathology.

Using SH-SY5Y cells as a model of neuronal hyperphosphorylation, we used a reverse-phase protein array (RPPA) analysis to demonstrate elevated levels of surface molecules specific to the hyperphosphorylative state (Fig 1C). Cell-SELEX capturing the differences between the surface of hyperphosphorylative cells and normal cells allowed the selection of phosphorothioate modified short DNA aptamers that bound with high affinity and specificity to hyperphosphorylative cells Fig 2 A. Having identified unique aptamers that bound such markers, we developed MR molecular imaging contrast agents that recognize the surface of cells in hyperphosphorylative state. We acknowledge that SH-SY5Y cells are not true neurons, they are a cell line originating in a neuroblastoma, a tumor of embryological neural crest origin. However, they can be induced to differentiate to a neuronal phenotype (as in the current work). While primary neuronal culture, or immortalized neuronal cells e.g. ReN-VM may offer alternative models of neurons, we have functionally tested the aptamer hits from our SELEX screen in a transgenic mouse model of tau deposition, and validated their performance, supporting our position that the choice of cell model was adequate to identify suitable markers of tau hyperphosphorylation.

PET is the leading modality for clinical molecular imaging, driven by its high contrast sensitivity, however, it suffers from poor spatial resolution on the order of 5-10mm, high cost; limited access to radioactive tracers, and radiation exposure. Nanoparticle-enhanced MR imaging overcomes all these obstacles, but historically has not achieved high enough sensitivity. We have previously demonstrated liposomal nanoparticles exhibiting large numbers of Gd chelates in the external bilayer leaflet, with hyper-T1 relaxive properties resulting in contrast sensitivity that rival nuclear imaging^45, 62^.

In P301S mice, the earliest reported histopathological studies are at age of 2.5months^63, 64^, and report no tau pathology. “Tau seeding” the cell-cell transfer of pathogenic tau aggregates has been reported using brain homogenates at 1.5 month of age^65^. We therefore chose P301S mice at 2 months of age for our studies when, tau seeding should be taking place, but frank tau pathology should be absent. The mice were injected with TauX nanoparticles targeted either by the Tau1 aptamer or the Tau3 aptamer. When imaged by T1-weighted MRI sequences, designed to optimize signal from the Gd chelate induced T1 relaxation caused by the liposomal-Gd nanoparticles, signal enhancement was observed in the cortex and hippocampus regions of the brain. Hyperphosphorylative conditions were confirmed by post-mortem IF staining with AT8 antibody that recognizes the S202 and T305 pTau species. Signal enhancement was not observed in non-transgenic mice, or in transgenic mice injected with untargeted nanoparticles, supporting the specificity of Tau1-or Tau3-bearing nanoparticle binding to target.

We have narrowed down the possible binding targets of the aptamers, and our data suggest that cell surface vimentin is a likely target. We have confirmed the specific presence of cell surface vimentin on the surface of SH-SY5Y cells in a hyperphosphorylative state, and on P301S mouse brain sections. Vimentin is an intermediate filament protein that undergoes constant assembly and/or remodeling and is usually associated with mesenchymal cells^66^. The assembly state of filaments is linked to their phosphorylation state, phosphorylation promotes disassembly^67^. Vimentin contains more than 35 phosphorylation sites targeted by multiple kinases and phosphatases allowing it to adjust IF dynamics dependent on its environment^68, 69^. Mechanical, chemical (toxins, hypoxia), and microbial stresses upregulate vimentin and its phosphorylation that allows cells to adjust their mechanical properties^70, 71^. The balance of different oligomeric forms influence dynamic cell processes including adhesion, migration, invasion including stress-induced signaling^72^. Vim IF’s (∼10nm) distributed throughout the cell by association with microtubules (tubulin, 24nm) regulating cell-migration, and microfilaments(actin, 7nm) regulating cell-contractility, form the cytoskeletal network and provide mechanical support for the plasma membrane where it contacts other cells or the extracellular matrix^73^. Interestingly, during the biological process epithelial to mesenchymal transition wherein non-motile, polar epithelial cells transform to motile invasive non-polar mesenchymal cells^74^, cells also undergo a cytoskeletal reorganization that includes changes in cell-membrane integrity, disassembly of junction proteins, increased stress-fiber formations, altered cell-surface protein expression. Changes in the localization of proteins is a hallmark of this pathologic process. Our observation that Vimentin is upregulated and translocated to the cell surface^75^ in the early stages of tau hyperphosphorylation suggests a possible role for EMT-related processes at the start of a slow progression towards AD pathology.

While positron emission tomography (PET) is the mainstay of molecular imaging, and exhibits remarkable sensitivity, there are several limitations posed by this methodology^62, 76^. Access to PET imaging is limited, even in the relatively well-served US, and is skewed towards high density urban centers. PET costs are very high due to the need for same-day radiosynthesis, and rapid decay of the isotopes. Longer half-life isotopes cause higher radiation exposure. This tradeoff between half-life and radiation exposure greatly limit the reach of PET to a wider patient population. Current PET tau tracers recognize the tau β-sheets in the PHF and NFT present in tauopathies^22^. This conformation is not unique to tau and the *in vivo* specificity is circumspect limiting its interpretation. Off-target binding of Flortaucipir, an approved tau PET agent, has been reported since it binds the MAO-B enzyme in the brain^77, 78^. Further, the vast majority of pathological tau is actually intracellular, posing a significant barrier to PET tracers that must navigate to the site of tau pathology, bind the target, and have all unbound tracer molecules cleared from the brain before the radioactive signal decays. Our choice of MRI as the detection modality is based on hyper-T1 relaxive properties of nanoparticles with surface conjugated Gd chelates, bringing detection sensitivity to the same range as nuclear imaging, and the MRI agent does not suffer from the rapid signal decay of PET agents, allowing plenty of time for unbound tracer to clear from the brain before imaging. Our choice of a cell surface surrogate marker of tau hyperphosphorylation avoids the need to bind an intracellular target. Finally, MRI imaging is already included in AD management and can be adjusted with agents such as TauX nanoparticles to constitute a highly sensitive and specific test for future tau pathology.

In summary, our work shows that the hyperphosphorylative conditions coinciding with the initiation of a decades long process culminating in AD result in markers manifested on the cell surface. By imaging their presence, our approach detects the presence of hyperphosphorylative cells indicative of a presymptomatic stage of AD. Coupled with early amyloid detection, this test could enable early positive diagnosis using the NIA-AA research framework.

## Materials and Methods

### Cell-lines

SH-SY5Y cells (ATCC, Manassas, VA, #CRL-2266™) were obtained from Dr. Jason Shohet’s lab at the Texas Children’s Hospital, Houston, immortalized human hepatocytes (THLE-3) were purchased from American Type Culture Collection (ATCC, Manassas, VA, # CRL-11233^TM^); both were cultured according to the ATCC instructions. ReN cell™ VM (#SCC008) cultured as per instruction using neural stem cell maintenance medium (#SCM005) and growth factors EGF (GF001) and bFGF(#GF005) all from Millipore Sigma, Burlington, MA.

### Differentiation

SH-SY5Y cells were exposed to 30 µM *all-trans*-Retionic acid (Sigma-Aldrich, St.Louis, MO, # R2625) in serum free cell medium for 10 days with medium change every alternate day. ReNcell VM were differentiated by the removal of growth factors from its culture medium for 10 days.

### Hyperphosphorylation

was induced in SH-SY5Y cells by addition of 30 nM okadaic acid (Sigma Aldrich, St.Louis, MO, # 459620) in growth medium with 30 µM RA for 24 hours. ReNcell VM were hyperphosphorylated using 100nM Quinolinic Acid (SigmaAldrich, St.Louis, MO, #P63204) in culture media for 24h.

### Synthesis of primers and TA DNA library

All primers and Cy5, and amine labelled selected aptamers were purchased from Integrated DNA Technologies (IDT, Coralville, IA). The ssDNA library used in Cell-SELEX contained a central randomized sequence of 30 nucleotides flanked by PCR primer regions to enable the PCR amplification of the sequence (5’-CGCTCGATAGATCGAGCTTCG-(N)_30_-GTCGATCACGCTCTAGAGCACTG-3’). The chemically synthesized DNA library was converted to a phosphorothioate modified library by PCR amplification using, dATP (αS), resulting in the DNA sequences where the 3’ phosphate of each residue is substituted with monothiophosphate groups, as described previously in detail ^32, 79^. The reverse primer was labeled with biotin to separate the sense strand from the antisense strand by streptavidin-coated sepharose beads (PureBiotech, Middlesex, NJ, # MSTR0510) for the next selection round. The concentration of the TA library was determined with a NanoDrop™ 2000 by measuring the UV absorbance at 260 nm.

### Cell-SELEX

The procedures of Cell-SELEX were performed according to protocol previously with some modification ^33, 80^.The initial ssDNA library of 150 pmole was dissolved in binding buffer with a total volume of 350 µl. It was denatured by heating at 95°C for 5 min, then renatured by rapid cooling on ice for 10 min. The treated SH-SY5Y cells at approximately 90% confluence in a 100-mm culture plate were washed twice with washing buffer and followed by incubating with the ssDNA library of 150 pmole for 2 hrs at 4°C. Following the incubation, for positive selections, the supernatant was discarded, and cells were washed three times with washing buffer to remove any unbound sequences. Then cells were scraped off and transferred to nuclease-free water, following another three times nuclease-free water washes. Cells in nuclease-free water was centrifuged at 300×g for 5 min. QIAamp DNA Mini and Blood Mini kit (Qiagen, Germantown, MD, # 51104) was introduced to elute cell membrane fraction. The cell membrane fraction was PCR-amplified to monitor the presence of cell binding efficacy at each cycle. For negative selections, the supernatant was simply pipetted out of the flask and processed for the next cycle of selection. The desired compartment were amplified by PCR, and used to prepare the TA for the next round of selection. Two different negative selections were involved. One was differentiated treatment only SH-SY5Y cells at cycles #12 and #13. Another was hepatocyte THLE-3 cells at cycles #20 and #21. A total of 26 cycles of Cell-SELEX were conducted, including two different types of negative selections mentioned above.

### Next-Generation Sequencing (NGS)

At the studied cycles, the membrane fractions were isolated and the recovered TA sequences were amplified by PCR. Equimolar quantities of the recovered TA sequences over the range were pooled together and sequenced by Next-Gen DNA sequencing using Ion318 chip (ThermoFisher, Waltham, MA). A four base sequence was introduced during PCR amplification to serve as unique “barcode” to distinguish between the studied cycles. Sequencing results were analyzed by the Aptalinger ^39^ that uses the markov model probability theory to find the optimal alignment of the sequences.

### Aptamer binding studies

were conducted with undifferentiated, differentiated and hyperphosphorylated SH SY5Y and ReN cell VM grown in 96-wells seeded at 10000 per well. The apparent dissociation constants (Kds) were measured by the equation Y=Bmax X/ (Kd+X), with GraphPad Prism 9, San Diego, CA, with a saturation binding experiment; cells were incubated with varying concentrations of Cy5-labeled aptamer in a 100 μl volume of binding buffer containing cells, incubated for 30minutes, washed twice and resuspended in 100 μl buffer and analyzed by Molecular probes microplate reader equipped with the appropriate excitation and emission filters. All data points were collected in triplicate.

### Immunocytochemistry

Eight-well glass plate was coated with a solution of 100 µg/ml Collagen Type I (Thermofisher Scientific, Waltham, MA, # A1064401) dissolved in 0.01N HCl and air dried, then PBS washed and air dried prior to seeding 20,000 SH-SY5Y cells per well. Aptamer staining at 100 nM was performed with live cells for 2h at 4⁰C in binding buffer and washed twice with washing buffer. Cells were then fixed by incubation for 15 min in 4% formaldehyde in PBS at room temperature. Non-specific binding was blocked with blocking buffer (G-Biosciences, St. Louis, MO, # 786195) for 1 hour and overnight incubation at 4°C with the rabbit pTau primary antibody (1:100) (Santa Cruz Biotechnology, #: sc-101815) was followed by washing with PBS, and 1h incubation with goat anti-rabbit IgG secondary antibody, Alexa Fluor 488 (Invitrogen, Carlsbad, CA, # A-11008) for 1 hour at room temperature. Cytoskeletal actin filaments were stained with Alexa Fluor 594 Phalloidin (Invitrogen, # A12381). The cells were covered with VECTASHIELD hardset mounting medium with DAPI (Vector Laboratories, Burlingame, CA, # H-1500) for 5 min at room temperature. Images were visualized under Olympus Fluoview FV1000 confocal microscopy.

### TauX nanoparticle synthesis

L-α-phosphatidylcholine, hydrogenated (Hydro Soy PC; HSPC) and Cholesterol were purchased from Lipoid Inc., Newark NJ, USA. 1,2-distearoyl-sn-glycero-3-phosphoethanolamine-N- [methoxy(polyethylene glycol)-2000] (DSPE-mPEG2000) was purchased from Corden Pharma, Liestahl, Switzerland. DSPE-PEG3400-COOH and Gd-DOTA-DSPE were synthesized *in house*, lis-rhodamine-DHPE from ThermoFisher Scientific. HSPC, Cholesterol, DSPE-PEG3400-COOH, DSPE-mPEG2000, Gd-DOTA-DSPE, lis-rhodamine-DHPE at molar proportions 31.4:40:0.5:3:25:0.1 were dissolved in ethanol to achieve a total concentration of 100 mM. For the non-targeted control stealth liposomes, carboxy terminated PEG was not included in the lipid mixture. The ethanolic solution of lipids was hydrated with 150mM saline solution at 65 °C for 30 minutes, allowing multilamellar liposomes to form. The mixture was then extruded in a 10ml Lipex extruder (Northern Lipids Inc., Burnaby, Canada) using a 400 nm polycarbonate track-etch polycarbonate filter (3 passes) followed by a 200 nm (3 passes) and finally 100nM filters. The suspension was then diafiltered using a MicroKros cross-flow diafiltration cartridge (*500 kDa* cutoff) from Repligen, Rancho Dominguez, CA, exchanging the external buffer for phosphate buffered saline (PBS, pH 7.2) for 15 volume exchanges. To form the aptamer conjugated liposomes, liposomes with lipid-PEG-COOH were reacted with amine terminated aptamers using carbodiimide chemistry. The carboxyl groups on the liposomes were activated with 5 mM EDC and 10 mM sulfo-NHS at pH∼6 for 5-10 minutes. The activated liposomes were then immediately reacted with the amine terminated aptamers and the pH was raised to ∼7.3 - 7.6 by titrating µl amounts of 5 N NaOH. The final concentration of aptamers used in reaction is ∼140 µM. The reaction was mixed at room temperature for 1 hr following which the reaction was carried out at 4 deg ^0^C overnight. The liposomes were then dialyzed against PBS to remove unconjugated aptamers using a 300 kDa dialysis membrane. The dialysate (external phase) was concentrated using 10 kD centrifugal separator and washed with PBS to remove residual EDC/s-NHS. The concentrated dialysate was analysed by NanoDrop Spectrophotometer (ThermoFisher Sci., Waltham, MA, USA) to determine unconjugated aptamer fraction, and estimate aptamer density per nanoparticle in TauX formulations. Inductively coupled plasma atomic emission spectroscopy (ICP-AES) was used to measure Gd and phosphorus concentrations of TauX formulations. The hydrodynamic diameter of liposomal nanoparticles in TauX formulations was determined using a dynamic light scattering instrument.

### Mice

All the procedures were performed with approval from Institutional Animal Care and Use Committee (IACUC) of Baylor College of Medicine. Mice were kept under a 12 h light/dark cycle, with food and water available ad libitum. PS19 mice from Jackson Laboratories (Bar Harbor, ME) B6; C3-Tg (Prnp-MAPT*P301S) PS19Vle/J Stock No: 008169 were used and experiments were conducted at the 2m of age. The transgenic (TG) mice develop neurofibrillary tangles by 5 months of age^64^. Age-matched non-transgenic wild type (WT) mice were used as controls.

### Magnetic Resonance Imaging (MRI)

MRI was performed on a 1T permanent magnet scanner (M7, Aspect Imaging, Shoham, Israel). Mice underwent pre-contrast baseline scans. Thereafter, mice were intravenously administered one of three nanoparticle MR contrast agents (TauT1, TauT3 or non-targeted control liposomes) via tail vein at a dose of 0.20 mmol Gd/kg of body weight. Delayed post-contrast MRI was performed 4 days after contrast agent injections. Pre-contrast and delayed post-contrast MR images were acquired using a T1-weighted spin echo (T1w-SE) sequence and a fast spin echo inversion recovery (FSE-IR) sequence with the following parameters: *<u>SE parameters</u>*: TR = 600 ms, TE = 11.5 ms, slice thickness = 1.2 mm, matrix = 192 × 192, FOV = 30 mm, slices = 16, NEX = 4; *<u>FSE-IR parameters</u>*: TR = 13500 ms, TE = 80 ms, TI = 2000 ms, slice thickness = 2.4 mm, matrix = 192 × 192, FOV = 30 mm, slices = 6, NEX = 6. Coil calibration, RF calibration, and shimming were performed at the beginning of study for each subject. The pre-contrast scans provide a baseline for calculation of signal enhancement from resulting post-contrast scans^46^. Two-standard deviations above the mean variation within WT control animals was used as the cutoff signal intensity for identifying tau positive animals. Six transgenic (TG) mice and six wild type mice (WT) were used for testing of each nanoparticle contrast agent formulation. Receiver operating characteristic (ROC) curves were generated on a six point ordinal scale by plotting the true positive fraction (TPF) against the false positive fraction (FPF) based on imaging-based identification of Tau-positive animals using the cutoff signal intensity and then comparing against histological confirmation of Tau pathology as a gold standard. A fitted curve was then generated against the empirical points plotted on the graphs. Qualitative and quantitative analysis of MRI images was performed in OsiriX (version 5.8.5, 64-bit, Pixmeo SARL, Geneva, Switzerland) and MATLAB (version 2015a, MathWorks, Natick, MA).

### Immunofluorescence

After the final MRI scan, the mice were euthanized and perfused extensively with 0.9% saline followed by 4% paraformaldehyde for 15 mins. The brains were then immersion-fixed in 4% formaldehyde for 48h at 4⁰C, transferred to 30% sucrose for cryoprotection and embedded in OCT. Phenotypic confirmation for the presence of phosphorylated tau and vimentin was done on 25µm thick brain sections. Antigen retrieval in pH=8.5 citrate buffer was executed in a 1200W GE microwave for 15mins. After 15 minutes of cooling 25µL of 1:50 dilution of primary p-tau antibody namely either AT8,AT100 or AT180 that recognize different p-tau species were incubated in a tray (RPI, Mt. Prospect, IL #248270) designed for microwave enhanced immunostaining procedures for 3min at power level 3. After a 2 min cooling, sections were washed with PBS and incubated for 3min with a 1:100 dilution of appropriate secondary antibody. DAPI staining proceeded after 2 mins of cooling and a PBS washing. ProGold Antifade (Invitrogen, Carlsbad, CA, # P36030) was used to mount slides which were visualized on Olympus Fluoview LV100. Scanning of whole sections was also conducted using a Biotek Cytation 5 slide scanning microscope. List of Antibodies – AT8 (#MN1020), Vimentin SP20 (#MA516409) both Thermo Fisher Scientific, Waltham, MA, Vimentin D21H3 (Cell Signaling Technology, Beverly, MA, #5741T,), Cell-surface vimentin (Abnova, Taipei City, Taiwan, #H00007431-M08J).

### Target Identification

The protein targets of Tau-1, Tau-3, Tau-4 and Tau-5 were identified by affinity-pull down using the selected aptamers as the capturing reagent followed by mass-spectroscopy. A scrambled DNA sequence, R2, was used as a control. The hyperphosphorylated SH-SY5Y cells, at 90-95% confluence were washed with cold PBS buffer and incubated with biotinylated selected aptamers with 25mmol/l each at 4°C in PBS, respectively. After 2 hours of gentle agitation, SH-SY5Y cells were cross-linked with 1% formaldehyde for 10 minutes at room temperature. The formaldehyde cross-linking was quenched with glycine. Cells were scraped from the plate, washed and lysed with lysing buffer (Thermofisher Scientific, # 87787) and treated with protease inhibitor mixture. The lysates were freeze-thawed for 30 minutes on ice and cleared by centrifuging at 10,000 ×g for 2 minutes at 4°C. To pull down the cross-linked proteins, equal amounts of cell lysate were incubated with prewashed streptavidin magnetic beads for 1 hour at room temperature with continuous rotation. Protein digestions were performed on the beads to isolate targeted proteins and processed for mass spectrometric analysis. Each sample was analyzed in triplicates. The raw data files were processed to generate a Mascot Generic Format with Mascot Distiller and searched against the SwissProt_2012_01 (Human) database using the licensed Mascot search engine v2.3.02 (Matrix Science, Boston, MA) run on an in-house server.

---

**Supplementary Table 1:**
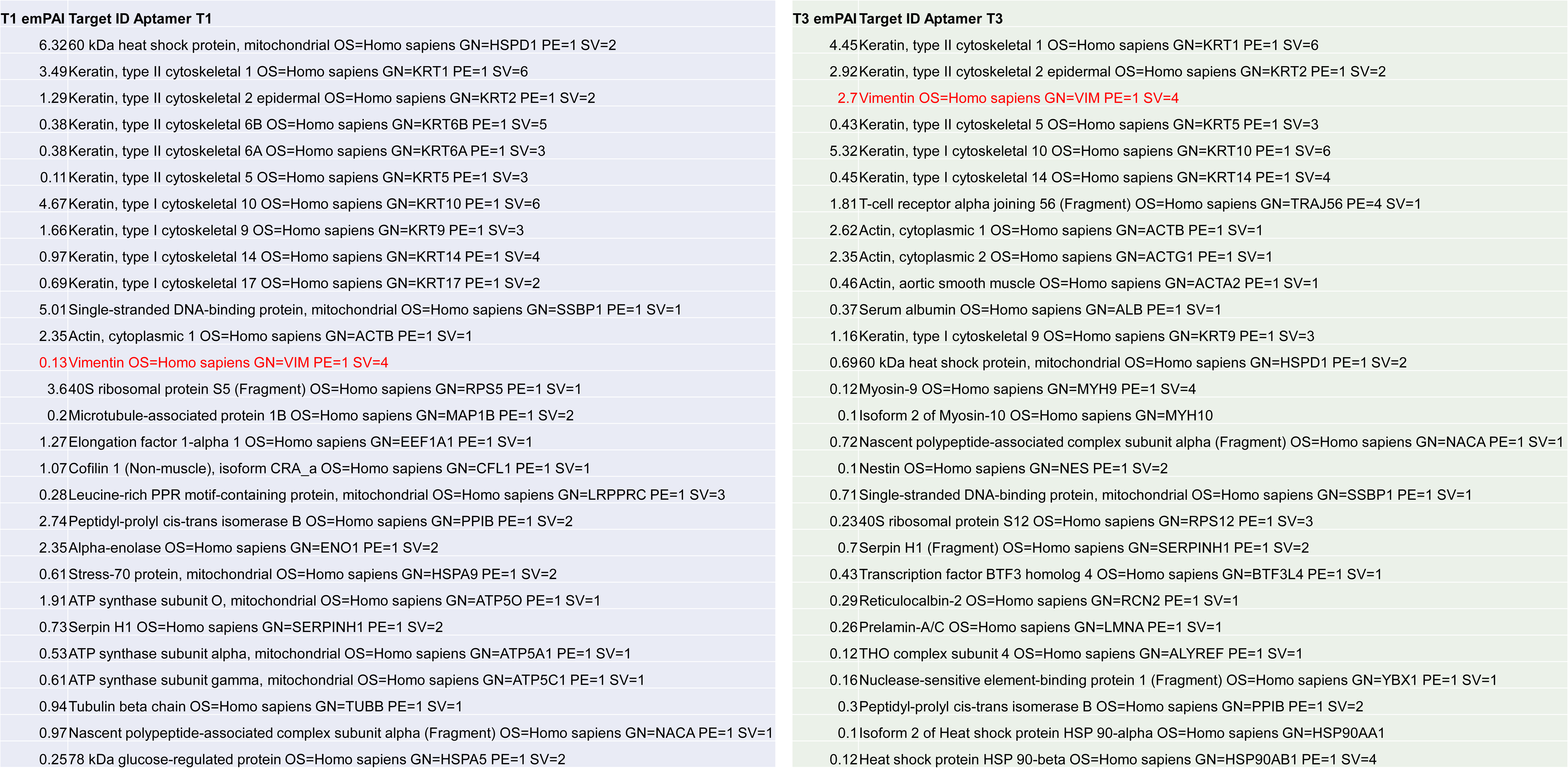
Target Identification using aptamer immunoprecipitation and mass spectrometry. Aptamer based immunoprepcipation was performed. Mass-Spectrometric analysis of the trypsin digested aptamer-bound complex was analyzed by LC/MS/MS on an LTQ-Orbitrap-XLmass spectrometer with an Nanoflex system. The raw data files were processed and searched against the SwissProt_2012_01 (Human) database using the Mascot search engine. Exponentially modified protein abundance index (emPAI) are reported in the table below that report protein content proportional to protein content in a mixture.

**Supplementary Figure S1:**
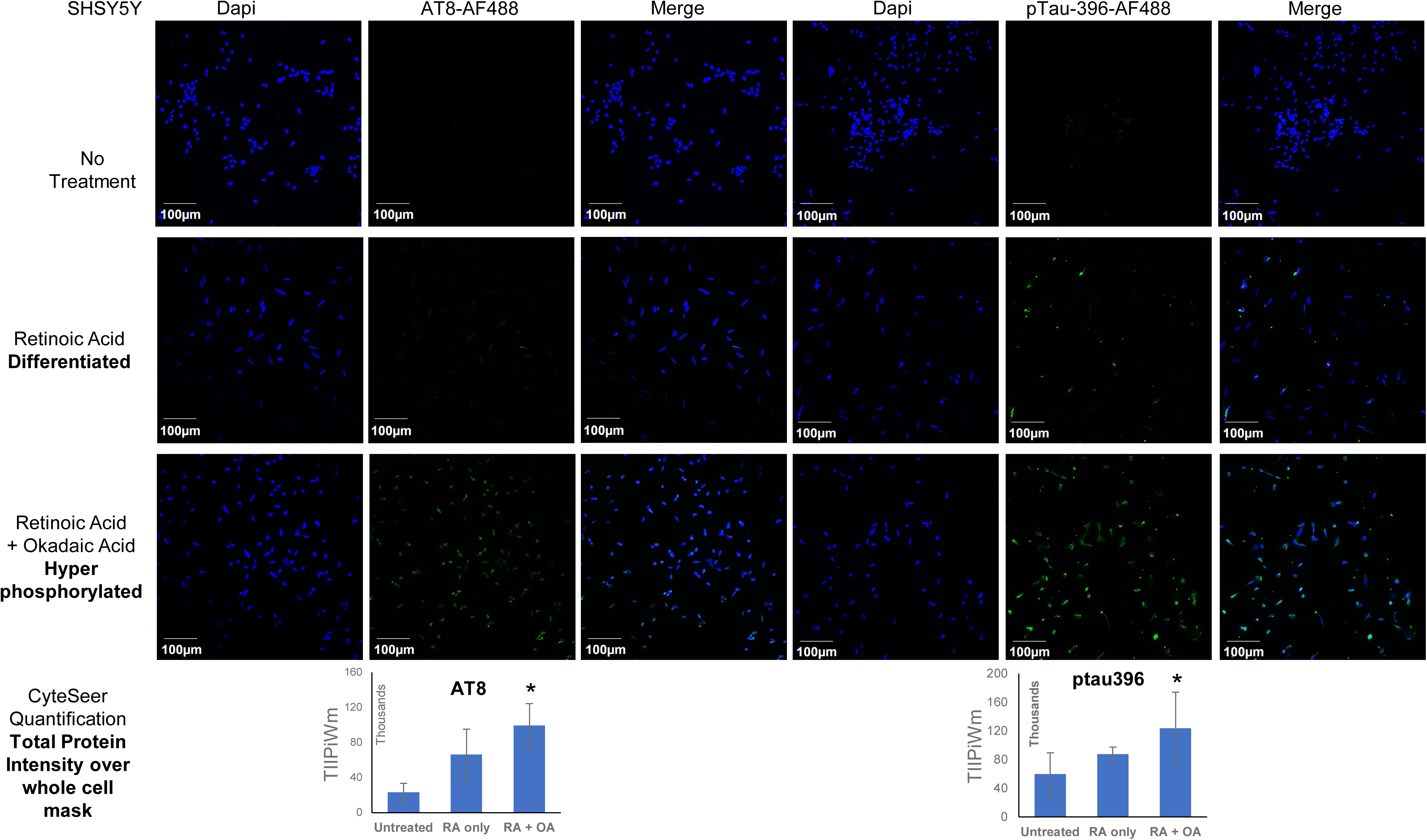
Hyperphosphorylative conditions. SHSY5Y cells were differentiated with 30µM Retinoic acid for 10 days. The differentiated cells were treated with 30nM Okadaic acid for 24hrs. Under these conditions Tau is hyperphosphorylated. We detected pTau Thr205/Ser202 and ptau 231 (an early phase of phosphorylation) and ptau Ser396 (a late stage of phosphorylation) stained using the antibodies AT8 and 5HCLC respectively. Quantification of the protein expression was conducted by using the automated image analysis software software CyteSeer that reports the total integrated protein intensity over the whole cell mask (TIIPiWm). A minimum of 5 different ROI’s with at least 50 cells were analysed. Each bar represents mean of means with SD. * p<0.05 compared to untreated control.

**Supplementary Figure S2:**
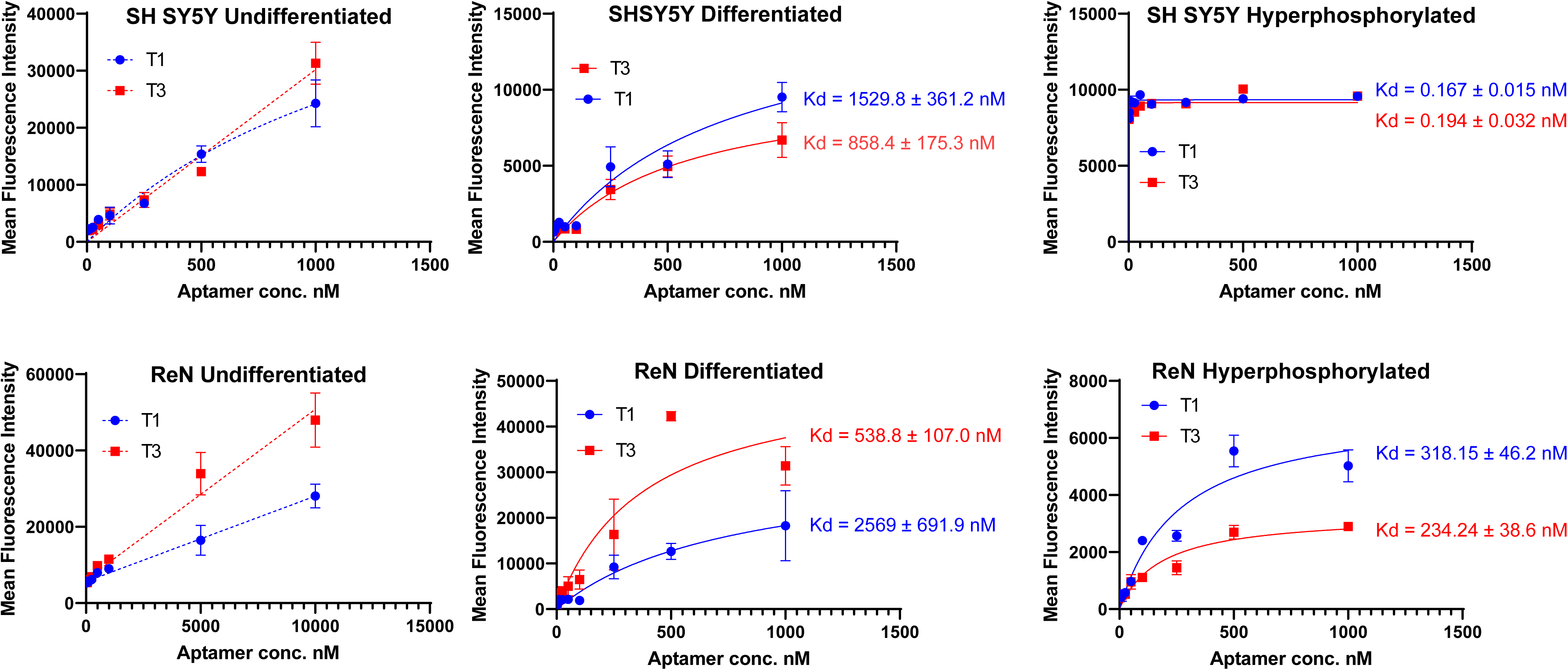
Apparent dissociation constants of Aptamer Tau1 and Tau3Target. SH SY5Y and ReN cells were grown in 96-well plates, Retinoic acid (30µM) was used for differentiation for 10 days. Okadaic acid (30nM) for SH SY5Y and Quinolinic acid (100nM) for ReN cells was used for 24 hrs to generate hyperphosphorylative conditions. Saturation binding curves were generated using Cy5 labelled Tau1 and Tau3 using a microplate fluorescent reader. The binding constant Kd apparent was calculated using the inbuilt non-linear regression module using the equation Y=Bmax*X/(Kd + X) in Graph Pad Prism. *\* error bars smaller than symbol not visualized*

## Notes

### Competing Interest Statement

The authors have declared no competing interest.

### Summary of Updates

Supplementary Figure 1 showing the presence of ptau Ser202/Thr 205 stained with antibody AT8 in addition to the ptau396. The caption to supplementary figure S1

